# Over four months of ethylene production: Unlocking the potential of solid-state photosynthetic cell factories

**DOI:** 10.1101/2025.09.12.675104

**Authors:** Sergey Kosourov, Vilja Siitonen, Gábor Tóth, Tuukka Levä, Tekla Tammelin, Pauli Kallio, Yagut Allahverdiyeva

## Abstract

This study demonstrates the feasibility of employing solid-state photosynthetic cell factories (solid-state PCFs) as a proof-of-concept platform for long-term ethylene production using sodium bicarbonate as the carbon source. Solid-state PCFs were constructed by entrapping *Synechocystis* sp. PCC 6803 (*efe* mutant, strain S5), specifically engineered for ethylene biosynthesis, within TEMPO-oxidized cellulose nanofiber (TCNF) matrices. Two distinct formulations were tested: (i) Ca^2+^-PVA-TCNF, in which TCNF was crosslinked with Ca^2+^ and polyvinyl alcohol to produce hydrogel films approximately 200 μm thick; and (ii) an all-polysaccharide-based Ca^2+^-MLG-TCNF formulation, in which TCNF was crosslinked with Ca^2+^ and mixed-linkage glucan. The latter films were fabricated using an osmotic dehydration approach, yielding mechanically robust, fully biodegradable structures with a thickness of approximately 2 mm. The integration of engineered cells with TCNF matrices created a biocatalytic system that improved the distribution of light, nutrients, and substrates to the cells, while facilitating ethylene separation, thereby supporting the fitness of immobilized cells and enhancing their metabolic performance. Using a custom-designed photobiofilm reactor optimized for semi-wet cultivation, the solid-state PCFs sustained ethylene production for over four months, representing the longest reported continuous ethylene production by cyanobacteria to date. Notably, the solid-state PCFs achieved up to a twofold increase in ethylene yield compared to the continuous-flow suspension culture. Importantly, the suspension-based system also represented the first demonstration of four-month ethylene production under continuous-flow operation. In addition, biodegradability assessments confirmed the environmental compatibility of the TCNF-based matrices, with the all-polysaccharide formulation being particularly advantageous due to its exclusively nature-based composition. Together, these results demonstrate the potential of solid-state PCFs as a scalable and sustainable platform for photosynthetic ethylene production.

## Introduction

Ethylene is an indispensable material in the chemical industry, serving as a precursor for a wide range of products, from plastics to antifreeze solutions.^1^ Additionally, it possesses high energy density (50.4 MJ kg^−1^), which is comparable to that of methane (55.5 MJ kg^−1^), making it an attractive candidate for use as a fuel.^2,3^ Currently, the industrial production of ethylene relies extensively on non-renewable petroleum resources, making the development of sustainable and renewable production pathways critically important.

Cyanobacteria provide an attractive carbon-neutral platform for the sustainable production of ethylene and other value-added chemicals, thanks to their ability to oxidize water and fix atmospheric CO_2_ through photosynthesis, using sunlight as the primary energy source. While cyanobacteria do not naturally produce ethylene, recent developments in molecular and synthetic biology have enabled their metabolic engineering and introduction of heterologous pathways for ethylene biosynthesis. A significant advancement in this field has been the expression of the ethylene-forming enzyme (Efe) from *Pseudomonas syringae* in various cyanobacterial hosts, including the model organism *Synechocystis* sp. PCC 6803.^2,4^ Introduction of the *efe* gene into cyanobacteria enables the cells to convert photosynthetically-fixed CO_2_ into ethylene *via* 2-oxoglutarate (2-OG), an intermediate of the tricarboxylic acid (TCA) cycle:

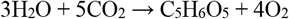

(photosynthetic CO_2_ fixation with formation of 2-OG),

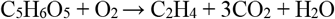

(conversion of 2-OG to ethylene by Efe).

It is important to note, however, that the above equations provide an oversimplified view of ethylene biosynthesis in the production host. In reality, the underlying interactions within the photoautotrophic metabolic networks are considerably more complex, involving the formation of side products and being influenced by competing carbon sinks that divert carbon flux away from ethylene biosynthesis.^5^ Moreover, 2-OG is not only a key intermediate of the TCA cycle but also a central hub linking carbon and nitrogen metabolism, serving as a precursor for amino acid biosynthesis and playing a regulatory role in nitrogen assimilation.^6^ This dual role further complicates the metabolic balance required for efficient ethylene production.

Optimization of ethylene production in engineered cyanobacteria has included the integration of codon-optimized versions of the *efe* gene under the control of various strong promoters and alternative ribosome binding site sequences.^7–9^ In parallel, engineering efforts have focused on rewiring the host’s central carbon metabolism to redirect carbon flux toward the TCA cycle, and more specifically, toward the formation of the Efe substrate 2-OG.^10,11^ These advances contribute to the development of cyanobacteria as photosynthetic cell factories (PCFs), a concept in which the production strain acts primarily as a biocatalyst, channelling energy and carbon directly into the synthesis of targeted chemical, while minimizing the generation of side-products.^12,13^ However, achieving the ideal performance for ethylene-producing PCFs remains challenging, as low productivity continues to be the primary bottleneck limiting the commercial viability of ethylene production by cyanobacteria.^14^

In practice, PCFs are typically deployed in photobioreactors as suspension cultures. However, suspension-based production platforms utilizing PCFs still face several inherent limitations. These include significant diversion of acquired energy and fixed carbon into biom ass growth rather than product formation. Other limitations are low light utilization efficiency due to self-shading and light scattering, excessive water consumption, the need for energy-intensive operations such as mixing and contamination control during cultivation, and costly downstream processes like biomass harvesting.^15,16^ Together, these challenges constrain the overall productivity and sustainability of current suspension-based production systems.

To achieve sustainable and economically viable production, fundamentally new technologies are necessary. Building upon a thin-layer immobilization approach originally developed to improve the photosynthetic production of H_2_,^17–20^ we recently introduced the concept of solid-state photosynthetic cell factories (solid-state PCFs). In this approach, specifically engineered photosynthetic cells are entrapped within a tailor-made, thin biobased polymeric matrix to enable sustainable and efficient production of solar chemicals and fuels.^21–23^ In solid-state PCFs, both the engineered cells and the functional properties of the matrix work synergistically to enhance production. The robust matrix restricts cell division and biomass accumulation, facilitates gas diffusion, and enables efficient product separation, while the engineered metabolic pathways in the cells channel energy and carbon fluxes toward the biosynthesis of the desired product. Furthermore, solid-state configurations allow for strategic light management by minimizing self-shading through the incorporation of structural elements within the matrix, such as heterogeneous cell aggregates,^24^ photosynthetic antenna gradients,^25,26^ modified photosynthetic antennae with broadened light absorption spectra,^27^ and integrated optical fibers. Notably, the introduction of a photosynthetic antenna gradient alone into the architecture of H_2_-producing biocatalysts dramatically improved light utilization efficiency, enabling light-to-H_2_ conversion efficiencies exceeding 4% at peak activity and maintaining over 2% throughout a continuous 16-day production period.^26^

In this research, we developed and applied ethylene-producing solid-state PCFs consisting of engineered cyanobacterial cells embedded within TEMPO-oxidized cellulose nanofiber (TCNF) matrices in a photobiofilm reactor (PhBFR). The PhBFR was designed for semi-wet cultivation of the biocatalyst to further enhance ethylene separation from the immobilization matrix. Building on this system, we demonstrated the successful application of solid-state cultivation technology, achieving continuous ethylene production for over four months, the longest reported duration for cyanobacterial ethylene production to date. This extended biocatalytic performance, combined with nearly twofold higher ethylene production yields compared to conventional suspension cultures, highlights the strong potential of solid-state PCF technology as a sustainable platform for CO_2_ capture and the photosynthetic production of value-added chemicals and fuels.

## Experimental

### Culture growth

The ethylene-producing strain *Synechocystis* sp. PCC 6803 pDF-lac2-S5-efe-CmR (hereafter referred to as *Synechocystis efe*) was used in the experiments.^8^ The cells were maintained in BG11 medium containing 20 mM HEPES-NaOH (pH 7.5) at 30°C, supplemented with 25 μg mL^−1^ spectinomycin and 10 μg mL^−1^ chloramphenicol. The culture flasks were placed in a versatile environmental chamber (MLR-351, Sanyo, Japan) under continuous shaking (120 rpm) on a rotary shaker in an atmosphere containing 1% CO_2_. Illumination was provided by continuous fluorescent light (Philips Master TL-D 36W/865) at approximately 35 μmol photons m^−2^ s^−1^ of photosynthetically active radiation (PAR).

Experimental cultures were grown without antibiotics and without agitation in 1 L flat round-neck Roux-type flasks (DURAN®, Schott, DWK Life Sciences GmbH) containing ∼800 mL of BG11 medium. These cultures were continuously bubbled with sterile filtered air supplemented with 1% CO_2_ (0.2 µm, Acro 37 TF, Gelman Sciences). Cells were harvested at OD_750_ of around 1 – 1.5 by centrifugation at 5000 g for 10 min (Avanti JXN-26, JA10, Beckman Coulter).

### Materials for cell immobilization

A water suspension of TEMPO-oxidized cellulose nanofibers (TCNF) with a final consistency of approximately 1 wt% was prepared as described previously.^21,23^ Polyvinyl alcohol (PVA) (Mowiol® 56-98, *M*_*W*_ ∼195 kg mol^−1^, DP 4300; Sigma-Aldrich) was dissolved in Milli-Q water at 90 °C in a water bath to prepare a 5 wt% solution. Low-viscosity barley mixed-linkage glucan (MLG) (*M*_*W*_ ∼179 kg mol^−1^; Megazyme, catalog number P-BGBL) was used to prepare a 1 wt% solution in Milli-Q water, as described previously.^28^ A poly(ethylene glycol) (PEG) solution (20 wt%) was prepared by dissolving solid PEG (*Mn* 35 kg mol^−1^) in Milli-Q water. All solutions and matrix formulations used for cell immobilization were sterilized by autoclaving.

### Fabrication of Ca^2+^-PVA-TCNF hydrogel films

For the preparation of Ca^2+^-PVA-TCNF hydrogel films, the collected *Synechocystis efe* pellet was resuspended in Milli-Q water at a ratio of 1 g (wet biomass) to 1 mL (Milli-Q water). The resulting suspension was then thoroughly mixed at a 1 : 1 ratio with a hydrogel formulation consisting of 1 wt% TCNF and 0.1 wt% PVA using a digital Ultra-Turrax homogenizer (IKA, Staufen, Germany) at 12000 rpm for 1 min. After mixing, air bubbles were removed by centrifugation at 3000 g for 5 min, and the final formulation was coated onto the surface of the Watman paper support using an automatic film applicator (TQC Sheen) and 200 µm-graded Mayer rods. The hydrogel matrix atop the paper was then stabilized by spraying 50 mM CaCl_2_ over the surface, resulting in a stable 7.5 cm x 16 cm (120 cm^2^) Ca^2+^-PVA-TCNF hydrogel film.

**Scheme 1.**
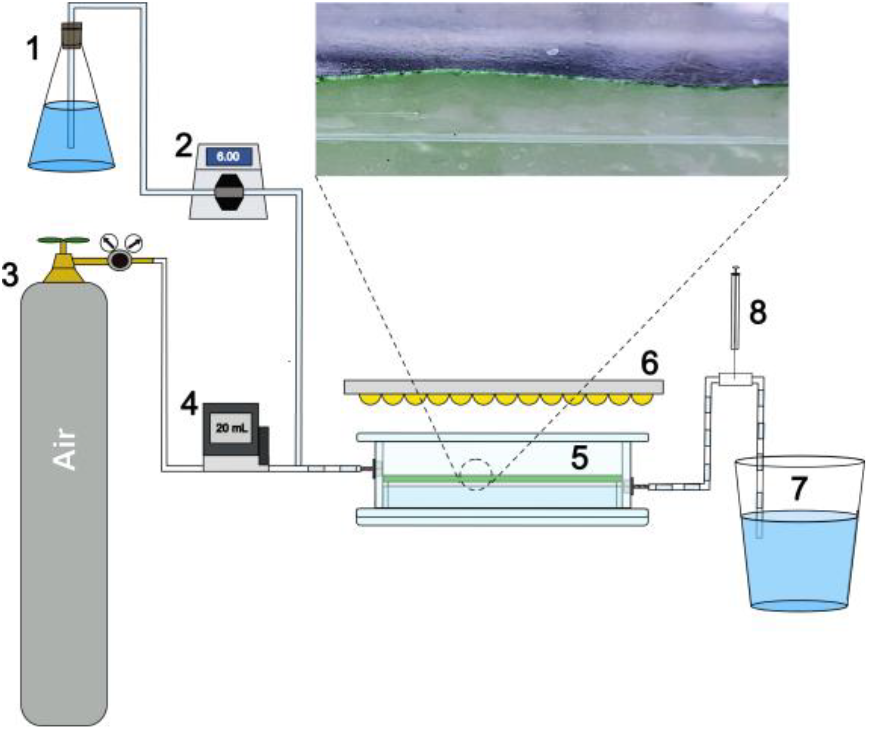
Schematic representation of the PhBFR setup for long-term ethylene production using solid-state photosynthetic cell factories. (1) Flask containing BG11 medium; (2) digital peristaltic pump; (3) compressed air cylinder; (4) digital mass flow controller; (5) PhBFR containing the photosynthetic thin-layer biocatalyst; (6) light panel; (7) flask for collecting discharged medium; (8) gas-tight syringe for gas sampling. The close-up shows the photosynthetic catalyst positioned atop the foam support inside the PhBFR.

The fabricated film was placed in the PhBFR on a water-conveying melamine foam support and left to adapt for 1 day in a closed PhBFR filled with 500 mL BG11 medium before the ethylene production phase. The PhBFR with the film was continuously illuminated from the top by fluorescence lamps (Philips Master TL-D 36W/865), providing approximately 40 μmol photons m^−2^ s^−1^ PAR.

### Fabrication of all-polysaccharide-based Ca^2+^-MLG-TCNF hydrogel film *via* osmotic dehydration

Bioinspired all-polysaccharide-based Ca^2+^-MLG-TCNF film with entrapped ethylene-producing *Synechocystis efe* cells was prepared using osmotic dehydration in a specialized casting frame, following the procedure described by Guccini et al.^29^ with modifications for the upscaled fabrication of rectangular (7.5 cm × 15 cm) films.

The film formulation was prepared by mixing 1 wt% TCNF and 0.1 wt% MLG with *Synechocystis efe* cell suspension (OD_750_ = 1 in Milli-Q water) at a 1:1 ratio using a digital Ultra-Turrax homogenizer. Air bubbles were removed by centrifugation at 3000 g for 5 min. The casting frame was 3D-printed using PLA and designed to hold a dialysis membrane (6–8 kDa cut-off), which was affixed to the bottom of the frame using super glue, forming an internal chamber for filling with 200 mL of the MLG-TCNF-cell formulation. The frame was placed into a second container with 200 mL of 20% PEG, ensuring full contact between the dialysis membrane and the PEG surface without air bubbles. The PEG solution in the second container was stirred with a magnetic bar on a stirring plate, and osmotic dehydration was carried out for 4 days. Following dehydration, the produced film was treated with 50 mM CaCl_2_, washed in Milli-Q water, and used directly for ethylene production without any further adaptation.

### Long-term ethylene production by immobilised and suspension cultures

The long-term ethylene production by the fabricated films was conducted in 1 L PhBFR with a 500 mL headspace under semi-wet cultivation conditions and continuous air and medium flows (Scheme 1). The PhBFR operated under sterile conditions throughout the experiment. The fabricated film was placed atop a water-conveying melamine foam support, which was partially submerged in 500 mL BG11 medium, ensuring that the film surface remained positioned a few millimetres above the liquid. This configuration allowed the entire film to remain exposed to the air atmosphere throughout the cultivation period.

The PhBFR with the film was continuously illuminated from the top by fluorescence lamps (Philips Master TL-D 36W/865), providing about 40 μmol photons m^−2^ s^−1^ PAR. The PhBFR system operated under continuous medium flow supplied by a digital peristaltic pump (#07525-20, MasterFlex) with a variable flow rate depending on the experimental setup and fixed air flow rate of 10 mL min^−1^ controlled by the high precision digital mass flow controller (Red-y Smart series, Vögtlin Instruments). The synthetic air (containing 21% O_2_ in N_2_, Woikoski Oy) was sterilized by passing through 0.2 µm filters (Acro 37 TF, Gelman Sciences). NaHCO_3_ (hereafter referred to as bicarbonate) was introduced into the BG11 medium at different concentrations depending on the experimental setup, serving as the carbon source. Ethylene production was initiated by the introduction of 1 mM IPTG in the PhBFR medium.

The experiment with suspension cultures was conducted in a similar PhBFR setup, with continuous mixing provided by a magnetic stirrer to prevent cell sedimentation and ensure homogeneity. The initial chlorophyll *a* (Chl *a*) concentration in the 500 mL suspension was adjusted to match the average total Chl *a* content in the fabricated films. The experimental conditions (i.e., medium and gas flow rates, bicarbonate concentration, temperature, and illumination) were identical for both setups, allowing a direct comparison of ethylene production between immobilized and suspension cultures.

The ethylene content in the PhBFR setup was monitored in the discharged gas flow using gas chromatography (GC, Clarus 580, PerkinElmer, Inc.). GC was equipped with Carboxen 1010 PLOT 30 m × 0.53 mm capillary column and a flame ionization detector set at 150°C with H_2_ and airflow fixed at 30 mL min^−1^ and 300 mL min^−1^, respectively. Argon (99.99%, Woikoski Oy) was used as a carrier gas at a flow rate of ∼9.8 mL min^−1^ (18 psi). The ethylene content was determined from 40 *μ*L gas samples collected using a gastight sample lock syringe (1705SL, Hamilton) several times per day. The average daily production yields were then calculated based on the air flow rate through the PhBFR and a calibration curve. The concentration of ethylene dissolved in the liquid phase was not considered due to its negligibly low solubility relative to its content in the gas phase. To express the production of ethylene in volumetric units, the amount of ethylene (mol) was converted to litres using the ideal gas law under the experimental conditions of 30°C and 1 atm (1 mol ≈ 24.87 L).

### Photosynthetic activity

The effective quantum yield of photosystem II [*Y(II)*] was measured using a PAM 2000 fluorometer (Walz, Germany) at several randomly selected spots across each film, both immediately after immobilization and at the end of the experiment. Prior to measurement, the film was dark-adapted for 10 minutes. A saturating light pulse (3000 μmol m^−2^ s^−1^ for 0.6 s) was applied on top of actinic light at 40 μmol m^−2^ s^−1^ to determine *Y(II)* at each spot.

### Chlorophyll determination

The Chl *a* concentration in the samples was determined spectrophotometrically as the absorbance at 665 nm minus 720 nm, following extraction of the cell pellets with 90% methanol, using the extinction coefficient reported by Lichtenthaler.^30^

### Sample preparation for enzymatic degradation

Dry TCNF films were prepared using the solvent-casting method, incorporating either PVA or MLG crosslinking additives, both of which have been shown to strengthen the TCNF network by bridging nanofibrils.^31,32^ PVA and MLG were added to their individual 0.5 wt% TCNF suspensions in amounts corresponding to 10% and 5% of the cellulose dry weight, respectively. The mixtures were homogenized using a high-performance dispersing instrument (Ultra-Turrax, IKA-Werke) at 13000 rpm for 2 min and degassed with a vacuum-assisted asymmetric centrifuge (SpeedMixer DAC 600, Synergy Devices Ltd.) operated at 1600 rpm and 100 mbar. Finally, the mixtures were cast into 8 cm Petri dishes and dried at 50% RH and 23°C for 4 days. The dried films were cut into approximately 1 cm x 1 cm pieces. The mean thicknesses of the dry PVA-TCNF and MLG-TCNF films were 22 µm and 26 µm, respectively.

Self-standing ∼1 wt% TCNF hydrogels, incorporating the same PVA and MLG additives as the dry films, were prepared following previously described methods.^23,28^ The materials were initially mixed at 0.5 wt% TCNF consistency, then cast onto a solid support and Ca^2+^-crosslinked to produce self-standing, low-solid-content hydrogels. After casting, the hydrogels were partially dewatered, then re-wetted by immersion in 25 ml of Milli-Q water and stored for 7 days at 4°C. Before enzymatic degradation, the hydrogels were cut into approximately 1 cm x 1 cm pieces. The Ca^2+^-PVA-TCNF and Ca^2+^-MLG-TCNF hydrogels had approximate thicknesses of 1 mm.

### Enzymatic degradation

The biodegradation assessment for the TCNF-based films and hydrogels was carried out with enzymatic degradation, following the method described by Leppänen et al.^33^ The enzyme mix was prepared according to Tenkanen et al.^34^ using commercially available enzymes: Gamanase, Novozyme 188 (Novozymes, Denmark), Econase and Ecopulp X-200 (AB Enzymes, Finland). The enzyme mix contained cellulase, mannanase, xylanase and β-glucosidase activities with a measured overall activity of 20 FPU/ml (filter paper unit/ml). The required volume of the enzyme mix was calculated according to Equation 1:

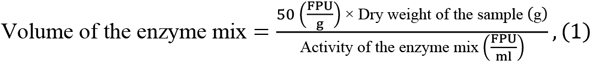

where a target enzyme activity of 50 FPU per gram of sample dry mass was used for the degradation. The experiments were conducted in 10 ml glass vials, where ∼1 cm^2^ film and hydrogel pieces with known dry matter contents were weighed. Then, sodium acetate buffer (0.5 M, pH 5) and MQ water were added to obtain a 0.7 wt% sample dry weight and 50 mM total ionic strength. Enzymatic degradation was initiated by adding the enzyme mix to the vials, which were then incubated at 40°C for 48 h with constant shaking. A reference sample containing only the enzyme mix was also prepared to account for background effects. After the incubation period, the samples were centrifuged at 3200 rpm for 10 min. The resulting supernatant, which contained the hydrolyzed sugar oligomers and enzymes, was boiled for 10 min to inactivate the enzymes. The precipitated enzymes were then separated by centrifugation and discarded. The reducing sugar content in the remaining supernatant was analysed using a dinitrosalicylic acid (DNS) reagent (1% DNS, 1.6% NaOH, and 30% potassium sodium tartrate), following the method described by Sumner and Noback.^35^ The degree of enzymatic hydrolysis for the samples was calculated with Equation 2:

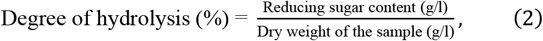

where the reducing sugar content in the supernatant was compared to the initial dry weight of the sample.

## Results and discussion

### Optimization of the long-term ethylene production by solid-state photosynthetic catalysts

To establish a stable and efficient system for long-term ethylene production, we entrapped *Synechocystis efe* cells within Ca^2+^-PVA-TCNF hydrogel films. The immobilization approach was implemented to enhance cell metabolic stability and prolong their biocatalytic activity, key factors in maintaining sustained photosynthetic productivity and long-term functionality over extended periods.^36–38^ The *efe* films were integrated into the PhBFR for semi-wet cultivation under continuous medium and air flows. This setup facilitated both ethylene release from the films and its monitoring.

In the first PhBFR setup, the medium contained 200 mM sodium bicarbonate, while its continuous flow rate varied between 5 mL h^−1^ and 30 mL h^−1^. The selection of 200 mM bicarbonate was based on previous findings demonstrating its effectiveness in sustaining ethylene production in both suspension and immobilized cultures.^39^ A two-month experiment revealed that a medium flow rate of 20 mL h^−1^, supplemented with 200 mM bicarbonate, was sufficient to sustain ethylene production (Fig. 1A). However, this setup led to the pronounced bleaching of the film after approximately 55 days of cultivation, accompanied by a gradual decline in ethylene production activity over time. Increasing the medium flow rate to 30 mL h^−1^ did not mitigate this degradation, suggesting that factors beyond nutrient availability may have contributed to the observed decline. Consequently, this setup yielded approximately 37.4 mmol (∼930 mL) of ethylene per m^2^ over 66 days of operation, with an average daily productivity of 0.57 mmol m^−2^ d^−1^ (∼14 mL m^−2^ d^−1^).

**Fig. 1.**
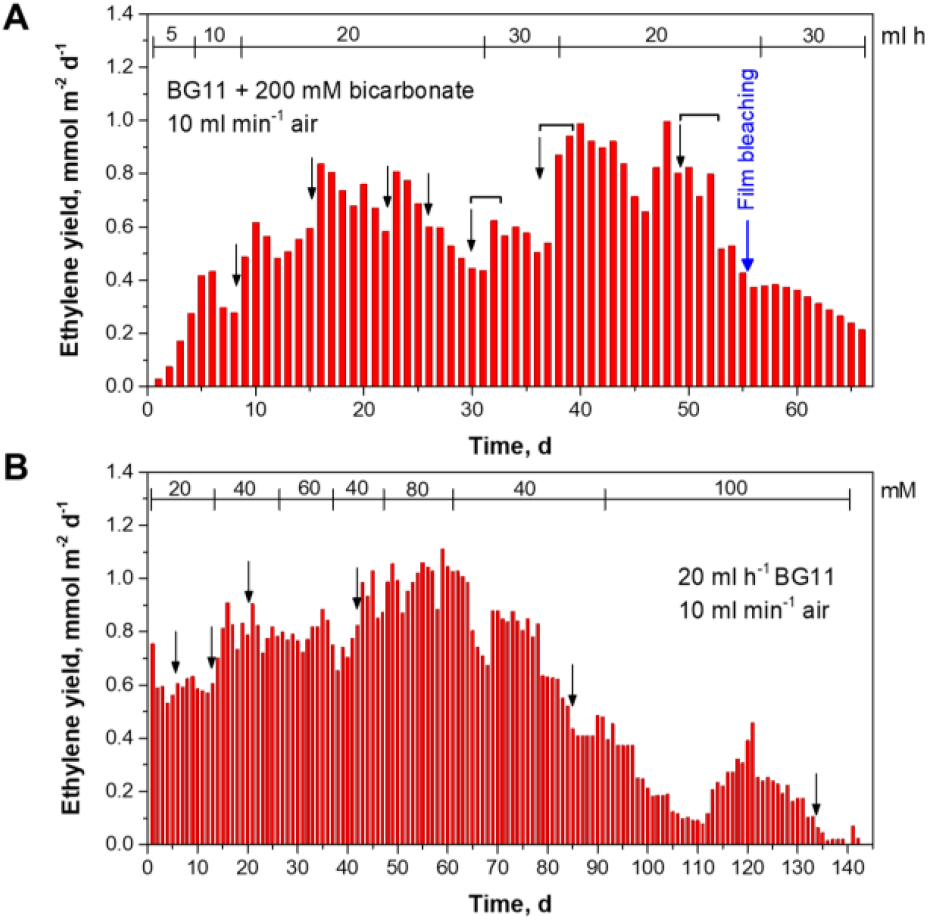
Continuous ethylene production by Ca^2+^-PVA-TCNF hydrogel films with entrapped Synechocystis efe cells under (A) varying BG11 medium flow rates with 200 mM sodium bicarbonate and (B) a fixed BG11 medium flow rate of 20 mL h^−1^ with different bicarbonate concentrations. Additions of 10 mM IPTG are indicated by arrows (introduced directly into the PhBFR) and arrows with brackets (introduced in the BG11 medium flask).

To further optimize the process, a second PhBFR setup was employed, where the flow rate was maintained at 20 mL h^−1^, and sodium bicarbonate concentration in the BG11 medium was varied from 20 to 100 mM. Under these conditions, the solid-state PCF catalyst sustained ethylene production for over 142 days, yielding approximately 80.4 mmol (∼2 L) of ethylene per m^2^ (Fig. 1B), with an average daily productivity similar to that observed in the first PhBFR (0.57 mmol m^−2^ d^−1^). A comparison of daily production yields between the two PhBFR setups (Fig. 1, panel A vs panel B) revealed that moderate bicarbonate loadings of 20 – 60 mM were sufficient to maintain daily ethylene production yields at levels comparable to those observed with 200 mM bicarbonate. Notably, introducing high bicarbonate concentrations during the later phase of cultivation (after approximately 90 days), when ethylene production activity began to decline, failed to restore ethylene production in immobilized cells (Fig. 1B). The data, obtained from the two PhBFR setups, suggest that carbon availability (Ci limitation) is not the primary factor responsible for the observed loss of ethylene-producing activity in the long-term process.

Another key finding from these experiments was that IPTG, an inducer of *efe* expression in *Synechocystis* cells, effectively triggered ethylene production during the early stages of the experiment. As cultivation progressed, immobilized cells no longer responded to IPTG addition (Fig. 1), indicating that further addition of the inducer did not affect ethylene production. Although the underlying causes of reduced inducer responsiveness in immobilized systems are not fully understood, our results suggest that this effect is more likely linked to the physiological state of the densely packed, non-growing cells rather than to limitations in IPTG diffusion. The initial responsiveness to IPTG during early cultivation confirms that the inducer was able to reach the cells effectively. However, the subsequent loss of responsiveness may also reflect properties of the expression system itself. The variant of the Lac promoter (P_A1lacO-1_) present in the *efe* strain used in this study has been reported to exhibit leaky expression in *Synechocystis*.^4,40^ According to Guerrero et al.,^4^ the accumulation of endogenous compounds such as allolactose in high-density cultures may interfere with LacI-mediated repression, potentially explaining the gradual loss of control over *efe* expression observed in our experiments. Therefore, stabilization of ethylene production over time may reflect a combination of leaky expression and sustained Efe protein stability in immobilized cells.

### Effect of bicarbonate loading on the long-term ethylene production by Ca^2+^-PVA-TCNF hydrogel films

Initial experiments described above revealed that moderate bicarbonate supplementation is sufficient to sustain ethylene production, suggesting that excessive bicarbonate addition may not provide further benefits. As an essential inorganic carbon source, HCO_3_^−^ supports CO_2_ fixation and maintains a sustained carbon flux for ethylene production.^11^ However, excessive supplementation can be detrimental, potentially inducing osmotic stress and disrupting the C/N balance in ethylene-producing cells.^41–43^ While 2-OG serves as the direct substrate for ethylene production in the *efe*-engineered strain, it also plays a broader role in cyanobacteria as a central metabolic hub linking carbon and nitrogen assimilation.^6,44^ Under conditions of limited nitrogen utilization, such as restricted growth due to cell entrapment within the matrix, nitrogen assimilation is reduced, potentially increasing intracellular 2-OG availability and thereby favouring ethylene biosynthesis. Such metabolic shifts, however, may also disturb the native carbon flux toward the GS-GOGAT cycle, adding complexity to the interpretation of production dynamics in solid-state PCFs. Beyond these physiological effects, the associated changes in the cultivation environment may also alter the physicochemical conditions within the matrix. For example, rapid bicarbonate consumption can locally raise pH, leading to carbonate precipitation, which may compromise matrix integrity and create localized carbon limitations.^22^

To evaluate the impact of higher bicarbonate concentrations on the solid-state PCFs, we operated three PhBFRs with Ca^2+^-PVA-TCNF films under identical medium flow rates, using BG11 medium supplemented with 20, 60, or 120 mM sodium bicarbonate. In all cases, IPTG was added only at the start of the experiment. As shown in Figure 2A, higher bicarbonate concentrations initially enhanced ethylene production yields, consistent with earlier short-term observations.^39^ However, during prolonged cultivation, high bicarbonate loadings led to a noticeably faster decline in daily production yields, accompanied by an earlier onset of visible film bleaching and reduced photochemical activity by the end of the experiment (Fig. 2B). To rule out genetic instability as a contributing factor, we confirmed that the decline in ethylene production was not caused by the loss or mutation of the plasmid carrying the *efe* gene in the immobilized cells under any of the tested experimental conditions (Supplementary Fig. 1). Consequently, total ethylene yields were 1.24, 0.72, and 0.68 L m^−2^ under 20-, 60-, and 120-mM bicarbonate, respectively. These results indicate that 20 mM bicarbonate is sufficient to sustain long-term biocatalytic activity, highlighting the importance of carefully controlling bicarbonate levels in the PhBFR to maintain cell viability and productivity in solid-state PCF catalysts. Moderate bicarbonate supplementation may also help maintain an optimal C/N balance, supporting stable 2-OG dynamics and efficient ethylene biosynthesis.

**Fig. 2.**
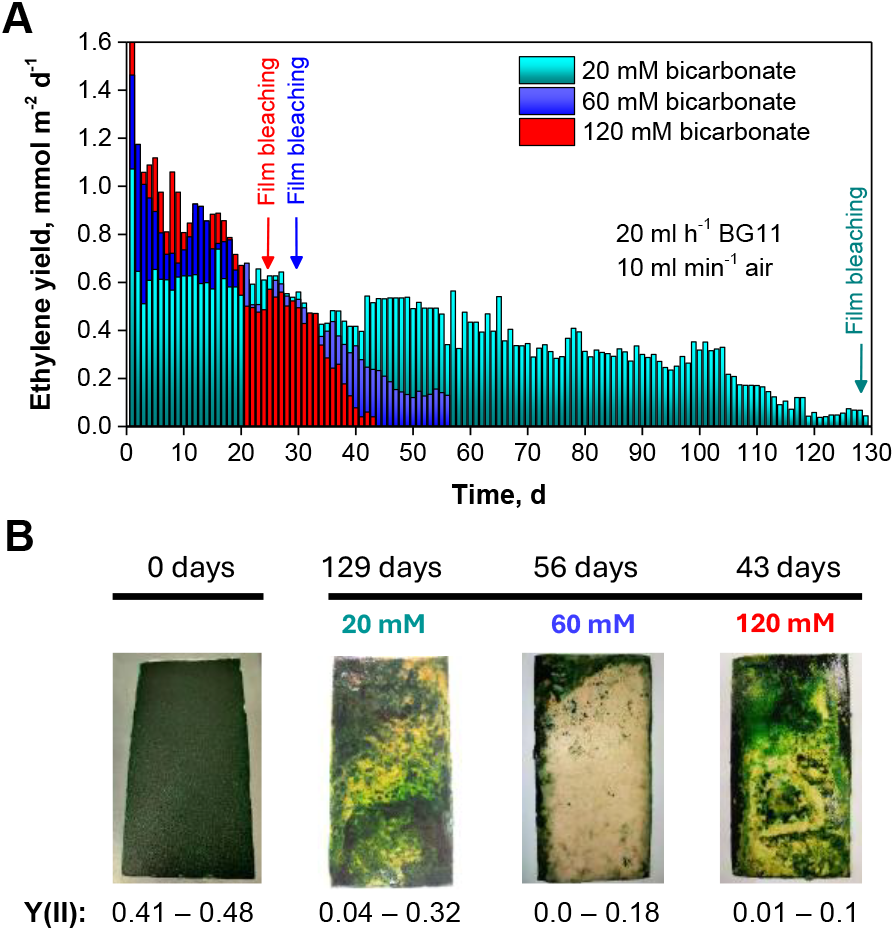
Effect of bicarbonate concentration on the long-term ethylene production and stability of Ca^2+^-PVA-TCNF hydrogel films. (A) Ethylene yield over time under continuous flow of BG11 medium (20 mL h^−1^) and constant air supply (10 mL min^−1^) for hydrogel films loaded with 20 mM (cyan), 60 mM (blue), and 120 mM (red) bicarbonate. Arrows indicate the onset of visible film bleaching. (B) Visual appearance and photochemical activity of the films at the beginning (0 days) and at the end of the experiment (43, 56, and 129 days at 120-, 60-, and 20-mM bicarbonate, respectively). Y(II) values represent the effective quantum yield of photosystem II, measured at several randomly selected spots across each film; the range indicates the minimum and maximum observed values.

While elevated bicarbonate concentrations negatively affect ethylene productivity, additional engineering limitations may also contribute to the long-term decline in biocatalytic performance. Structural degradation of the immobilization matrix can result in cell loss, while diffusion limitations restrict gas, nutrient, and product exchange, causing localized substrate depletion, excessive O_2_ accumulation, and matrix heterogeneity.^45–47^ The presence of matrix heterogeneity is evident in the photographs taken at the end of the ethylene production experiment, where uneven film bleaching and discoloration reflect variations in biocatalyst condition (Fig. 2B). These factors destabilize the microenvironment within the matrix, impairing cellular activity and further reducing ethylene yields.

The experiments thus demonstrated that optimizing both bicarbonate levels and the physicochemical properties of the immobilization matrix is essential for sustaining high biocatalytic efficiency in PhBFRs. Further engineering efforts should focus on increasing matrix porosity to enhance mass transfer and maintain a stable, uniform biocatalytic environment over extended operation periods.

### Comparison of the hydrogel films with conventional suspension cultivation

To confirm the advantages of the immobilization approach over the traditional suspension-based production system, we operated the PhBFR loaded with a suspension *efe* culture. The total cell density was adjusted to match that of the immobilized films, normalized based on Chl *a* content. The reactor was supplied with BG11 medium containing 20 mM NaHCO_3_ at a flow rate of 20 mL h^−1^. To prevent cell sedimentation, the suspension culture in the PhBFR was continuously and vigorously mixed using magnetic stirring throughout the experiment. This experiment represents one of the first demonstrations of long-term ethylene production by a photosynthetic suspension culture under controlled flow conditions.

The suspension culture cumulatively produced 1.8-fold less ethylene than the immobilized system under the same operational conditions, yielding 26.6 mmol m^−2^ (∼0.66 L m^−2^) of ethylene over 129 days (Fig. 3), compared to 50 mmol m^−2^ (1.24 L m^-2^) produced by the Ca^2+^-PVA-TCNF hydrogel film over the same period (Fig. 4). The average daily productivities were approximately 5 mL m^−2^ d^−1^ for the suspension culture and 10 mL m^−2^ d^−1^ for the film. This performance gap aligns with previous findings showing that free-living cyanobacterial cultures typically yield less ethylene over prolonged cultivation, primarily due to greater diversion of energy toward biomass accumulation rather than product synthesis, and potentially lower availability of 2-OG for ethylene biosynthesis resulting from increased metabolic demand in actively dividing cells.^5,48^ In our system, the performance of suspension cultures was further affected by cell washout under continuous medium flow conditions. Indeed, the PhBFR with the suspension culture maintained a relatively low cell density (below 1 mg Chl *a* per PhBFR) for the majority of the cultivation period (Fig. 3), which likely minimized the contribution of self-shading effect.

**Fig. 3.**
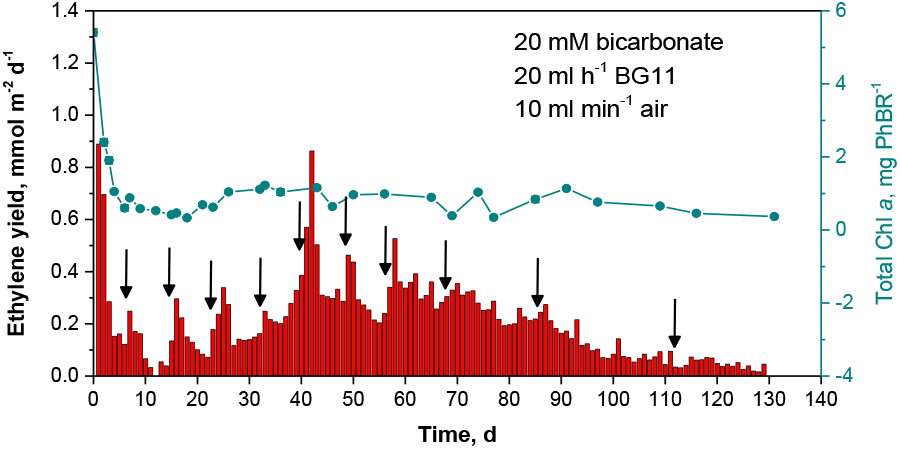
Daily ethylene production yields by the suspension of *Synechocystis efe* cells and the change of Chl a content in the PhBFR throughout long-term cultivation. Additions of 10 mM IPTG in the PhBFR are indicated by arrows.

**Fig. 4.**
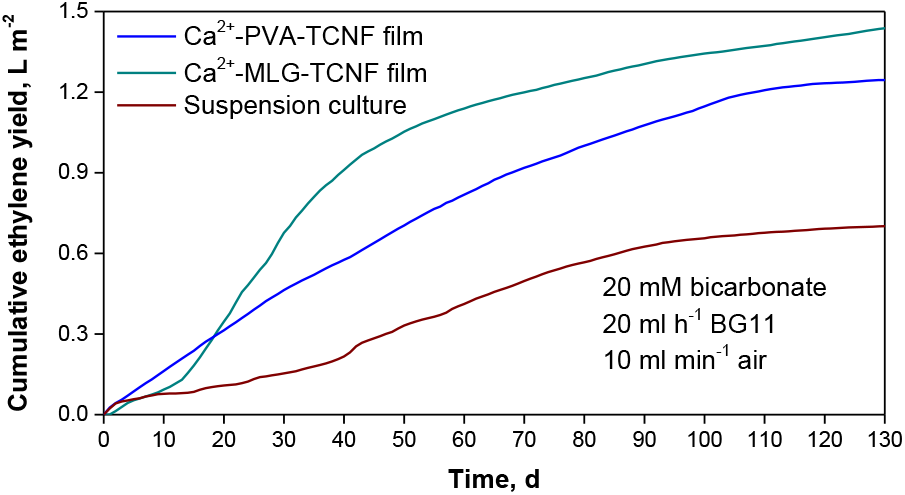
Comparison of cumulative ethylene production yields by suspension culture, Ca^2+^-PVA-TCNF and Ca^2+^-MLG-TCNF hydrogel films.

Unlike the immobilized system, the suspension culture required periodic additions of 10 mM IPTG to maintain *efe* expression. This was necessary due to the continuous growth of the culture and dilution effects, which gradually reduced intracellular inducer levels over time. Additionally, the requirement for constant mixing to prevent cell sedimentation increased the overall maintenance demands, making suspension-based ethylene production less efficient in long-term operation. These findings support the advantages of immobilization, where enhanced cell retention contributes to improved ethylene yields and long-term productivity. At the same time, reduced maintenance requirements offer additional benefits from an operational and economic perspective.

### Bioinspired formulation for long-term ethylene production

In the final set of ethylene production experiments, we selected an all-polysaccharide-based Ca^2+^-MLG-TCNF matrix formulation as the most promising candidate for long-term application.^32^ This choice was based on its excellent performance in entrapping *Synechocystis efe* cells in short-term ethylene production.^28^ Unlike the previous study by Virkkala and co-authors,^28^ which relied on conventional atmospheric drying, we adopted an osmotic dehydration technique to prepare the films. This approach enabled the formation of a porosity gradient across the matrix volume, resulting in more uniform dehydration and enhanced mass transport properties compared to standard drying methods.^29^ In addition, the MLG-TCNF formulation produced films with excellent mechanical stability,^28^ a critical factor for maintaining structural integrity and function during extended cultivation.

As shown in Fig. 4, the all-polysaccharide-based film outperformed the Ca^2+^-PVA-TCNF formulation in total ethylene production yield under identical cultivation conditions. However, it is important to note that the osmotic dehydration approach produced a significantly thicker film, approximately 10 times thicker than the Ca^2+^-PVA-TCNF matrix. When normalized to catalyst volume, this led to a markedly lower space-time yield of 0.22 mmol ethylene L_cat_ ^-1^ d^-1^ for the 2 mm Ca^2+^-MLG-TCNF film, compared to 1.9 mmol ethylene L_cat_ ^-1^ d^-1^ for the 200 *μ*m Ca^2+^-PVA-TCNF film. The reduced volumetric productivity of the thicker MLG-TCNF film is most likely attributed to pronounced self-shading and light limitation within the matrix. While thinner films produced *via* optimized osmotic dehydration could help mitigate this issue, they may come at the expense of total productivity due to reduced cell loading. Alternatively, more advanced architectural designs of the biocatalyst could be explored. These include cell clustering for better light distribution,^24^ gradual truncation of photosynthetic antennae to reduce ineffective light absorption within the matrix^26^ and/or the incorporation of novel materials to enhance internal light distribution throughout the film. Such strategies could help balance light availability, cell density, and overall productivity in future immobilized systems.

Nonetheless, both immobilized systems dramatically outperformed the suspension culture in terms of space-time yield, which remained around 0.007 mmol ethylene L_cat_ ^-1^ d^-1^. These results demonstrate the clear advantage of the solid-state PCF architecture for long-term ethylene production, combining improved productivity with reduced operational demands.

### Biodegradation assessment for TCNF-based dry films and hydrogels

To complement the long-term ethylene production experiments, we assessed the biodegradability of the developed TCNF-based immobilization matrices. Both self-standing hydrogels and completely dried films (Fig. 5A) were subjected to rapid laboratory-scale enzymatic degradation experiments. After 48 hours of incubation, the dry PVA-TCNF and MLG-TCNF films had completely disintegrated, while the corresponding Ca^2+^-crosslinked hydrogels had liquefied. The reducing sugar content analysis revealed that 65 – 70% of the films’ and 35 – 40% of the hydrogels’ dry matter had hydrolyzed to soluble sugar oligomers (Fig. 5B). Slightly higher degradation was observed for the all-polysaccharide MLG-crosslinked materials compared to those containing synthetic PVA.

**Fig. 5.**
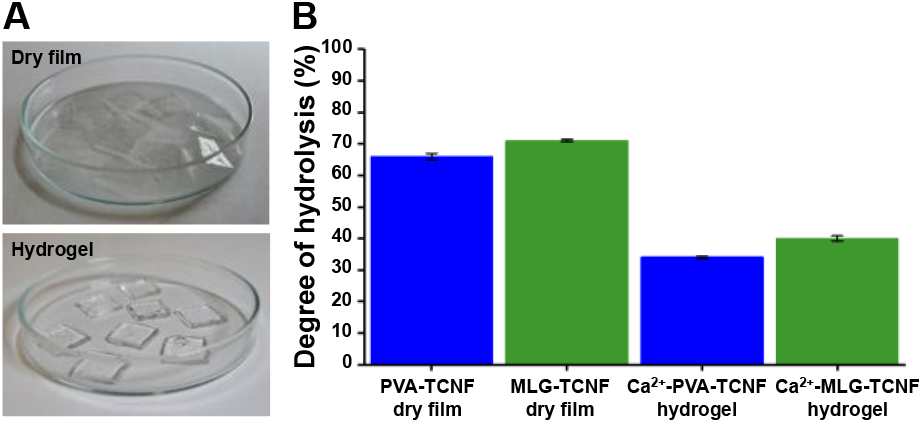
Photographs of the TCNF-based dry films and hydrogels prior to enzymatic degradation experiments (A). The percentage of TCNF-based dry films’ and hydrogels’ dry matter hydrolyzed to sugars after 48 h enzymatic degradation (B). Samples were prepared with the crosslinking additives PVA (blue), MLG (green) and Ca^2+^-ions (only in hydrogels). Enzymatic degradation was conducted in duplicate for each sample type and degree of hydrolysis determined twice from each duplicate.

A more pronounced difference was noted between the dry films and hydrogels: regardless of crosslinking additive, the dry films degraded more extensively. This is likely explained by improved enzyme accessibility in the case of thin film samples, whereas the bulkier hydrogel structure limited enzymatic hydrolysis due to a combination of slow diffusion and a lower effective substrate concentration. Nevertheless, given more time, the TCNF-based hydrogels would likely degrade as thoroughly as the dry films. Previous studies have indeed demonstrated excellent enzymatic degradation of TCNF samples,^49^ as well as extensive microbial mineralization under both aerobic and anaerobic conditions.^50^ In the latter case, TCNF exhibited near-complete biodegradation comparable to that of native cellulose nanofibers, although at a reduced rate. This decrease in degradation rate has been shown to correlate well with the increasing degree of substitution in the cellulose derivatives, which can hinder microbial or enzymatic access to the cellulose backbone.^33^ Despite such variations in degradation kinetics, all native and chemically modified cellulose-based films that exhibited partial degradation in 48-hour enzymatic tests were found to fully disintegrate in prolonged, pilot-scale composting experiments.^33^

Taken together, our enzymatic degradation experiments confirm the high degradability of TCNF-based immobilization matrices, regardless of whether MLG or PVA was used as the crosslinking additive. However, it is important to note that the degradation of PVA-containing materials could potentially contribute to microplastic formation.^51^ This concern further emphasizes the importance of advancing all-polysaccharide immobilization matrices that use MLG or other bio-based crosslinkers, as they offer a fully biodegradable and environmentally safer alternative to PVA-based materials for long-term applications.

## Conclusions

This study demonstrates the considerable potential of solid-state PCFs as a platform for long-term light-driven ethylene production, based on cyanobacterial cells entrapped within thin nanocellulose matrices. Specifically, the engineered *Synechocystis efe* cells were introduced within Ca^2+^-PVA-TCNF and Ca^2+^-MLG-TCNF formulations, forming stable biocatalysts that sustained ethylene biosynthesis under ambient conditions for up to four months without reinduction or catalyst replacement. This contrasts sharply with state-of-the-art electrochemical CO_2_-to-ethylene devices, which typically degrade within 14 to 200 h due to catalyst deactivation or electrolyte breakdown,^52,53^ whereas direct hydrogenation of CO_2_ requires high-temperature and high-pressure conditions.^54,55^

In addition, the solid-state PCFs significantly outperformed the suspension PCFs in terms of productivity, despite being operated under identical conditions and with the same initial cell density. It is worth noting that the suspension system itself represented the first demonstration of long-term ethylene production under continuous-flow conditions. Optimization studies also revealed that, in continuous-flow cultivation, moderate bicarbonate concentrations (20–60 mM) were sufficient as an inorganic carbon source to maintain prolonged ethylene production while avoiding the negative effects observed at higher bicarbonate loadings.

The use of an osmotic dehydration approach to fabricate mechanically stable biocatalysts introduced an important innovation, improving both the structural integrity and mass transport properties of the films, as confirmed in long-term operation. Biodegradability assessments further demonstrated the environmental sustainability of these TCNF-based matrices, with all-polysaccharide formulations showing particularly favourable enzymatic degradation profiles compared to matrices crosslinked with synthetic polymers such as PVA. Altogether, the findings presented here underline the viability of solid-state PCF technology as a robust, scalable, and environmentally sustainable platform for photosynthetic chemical production.

## Author contributions

Conceptualization and methodology – SK, YA; investigation of long-term ethylene production – SK, VS, GT, PK, YA; investigation of biodegradation – TL, TT; data analysis – SK, TL, GT, VS; resources – YA, PK, TT; funding acquisition – YA, TT; writing (original draft on long-term ethylene production) – SK; writing (original draft on biodegradation) – TL; writing (review & editing) – all authors.

## Acknowledgements

This work was financially supported by the EU FET Open Project FuturoLEAF under grant agreement no. 899576, and the Jane and Aatos Erkko Foundation (PhotoFactory project). SK and YA thank Dr. Dan Enke from Cyano Biotech (now Simris Biologics, Germany) for providing the mass flow controllers, and Tiia Siivola for her assistance with ethylene production in one of the photobioreactor setups. TL and TT thank Elisa Spönla and Riitta Alander (VTT Technical Research Centre of Finland) for their help with the enzymatic degradation experiments.

## References

1 R. A. Sheldon, Phil. Trans. R. Soc. A., 2024, 382, 20230259.

2 J. Ungerer, L. Tao, M. Davis, M. Ghirardi, P.-C. Maness and J. Yu, Energy Environ. Sci., 2012, 5, 8998–9006.

3 K. Ogura, Journal of CO2 Utilization, 2013, 1, 43–49.

4 F. Guerrero, V. Carbonell, M. Cossu, D. Correddu and P. R. Jones, PLoS ONE, 2012, 7, e50470.

5 P. Kallio, A. Kugler, S. Pyytövaara, K. Stensjö, Y. Allahverdiyeva, X. Gao, P. Lindblad and P. Lindberg, Physiologia Plantarum, 2021, 173, 579–590.

6 K. Forchhammer and K. A. Selim, FEMS Microbiology Reviews, 2020, 44, 33–53.

7 V. P. Veetil, S. A. Angermayr and K. J. Hellingwerf, Microb Cell Fact, 2017, 16, 34.

8 K. Thiel, E. Mulaku, H. Dandapani, C. Nagy, E.-M. Aro and P. Kallio, Microb Cell Fact, 2018, 17, 34.

9 B. Wang, C. Eckert, P.-C. Maness and J. Yu, ACS Synth. Biol., 2018, 7, 276–286.

10 S. Lynch, C. Eckert, J. Yu, R. Gill and P.-C. Maness, Biotechnol Biofuels, 2016, 9, 3.

11 C. Durall, P. Lindberg, J. Yu and P. Lindblad, Biotechnology for Biofuels, 2020, 13, 16.

12 G. Luan and X. Lu, Biotechnology Advances, 2018, 36, 430–442.

13 A. Melis, D. A. Hidalgo Martinez and N. Betterle, Photosynth Res, 2024, 162, 459–471.

14 J. N. Markham, L. Tao, R. Davis, N. Voulis, L. T. Angenent, J. Ungerer and J. Yu, Green Chem., 2016, 18, 6266–6281. 15

15 G. Torzillo, B. Pushparaj, J. Masojidek and A. Vonshak, Biotechnology and Bioprocess Engineering, 2003, 8, 338–348.

16 L. Novoveská, S. L. Nielsen, O. T. Eroldoğan, B. Z. Haznedaroglu, B. Rinkevich, S. Fazi, J. Robbens, M. Vasquez and H. Einarsson, Marine Drugs, 2023, 21, 445.

17 J. L. Gosse, B. J. Engel, F. E. Rey, C. S. Harwood, L. E. Scriven and M. C. Flickinger, Biotechnology Progress, 2007, 23, 124–130.

18 S. N. Kosourov and M. Seibert, Biotechnology and Bioengineering, 2009, 102, 50–58.

19 J. L. Gosse, B. J. Engel, J. C.-H. Hui, C. S. Harwood and M. C. Flickinger, Biotechnology progress, 2010, 26, 907–918.

20 H. Leino, S. N. Kosourov, L. Saari, K. Sivonen, A. A. Tsygankov, E. M. Aro and Y. Allahverdiyeva, International Journal of Hydrogen Energy, 2012, 37, 151–161.

21 M. Jämsä, S. Kosourov, V. Rissanen, M. Hakalahti, J. Pere, J. A. Ketoja, T. Tammelin and Y. Allahverdiyeva, Journal of Materials Chemistry A, 2018, 6, 5825–5835.

22 V. Rissanen, S. Vajravel, S. Kosourov, S. Arola, E. Kontturi, Y. Allahverdiyeva and T. Tammelin, Green Chem., 2021, 23, 3715–3724.

23 T. Levä, V. Rissanen, L. Nikkanen, V. Siitonen, M. Heilala, J. Phiri, T. C. Maloney, S. Kosourov, Y. Allahverdiyeva, M. Mäkelä and T. Tammelin, Biomacromolecules, 2023, 24, 3484–3497.

24 S. T. Chua, A. Smith, S. Murthy, M. Murace, H. Yang, L. Schertel, M. Kühl, P. Cicuta, A. G. Smith, D. Wangpraseurt and S. Vignolini, Proc. Natl. Acad. Sci. U.S.A., 2024, 121, e2316206121.

25 S. N. Kosourov, M. He, Y. Allahverdiyeva and M. Seibert, in Microalgal Hydrogen Production: Achievements and Perspectives, The Royal Society of Chemistry, 2018, pp. 355–384.

26 S. Kosourov, T. Tammelin and Y. Allahverdiyeva, Energy Environ. Sci., 2025, 18, 937–947.

27 R. Croce and van A. Herbert, Nature Chemical Biology, 2014, 10, 492–501.

28 T. Virkkala, S. Kosourov, V. Rissanen, V. Siitonen, S. Arola, Y. Allahverdiyeva and T. Tammelin, J. Mater. Chem. B, 2023, 11, 8788–8803.

29 V. Guccini, J. Phiri, J. Trifol, V. Rissanen, S. M. Mousavi, J. Vapaavuori, T. Tammelin, T. Maloney and E. Kontturi, ACS Appl. Polym. Mater., 2022, 4, 24–28.

30 H. K. Lichtenthaler, Methods in Enzymology, 1987, 148, 350–382.

31 M. Hakalahti, A. Salminen, J. Seppälä, T. Tammelin and T. Hänninen, Carbohydrate Polymers, 2015, 126, 78–82.

32 S. Arola, M. Ansari, A. Oksanen, E. Retulainen, S. G. Hatzikiriakos and H. Brumer, Soft Matter, 2018, 14, 9393–9401.

33 I. Leppänen, M. Vikman, A. Harlin and H. Orelma, J Polym Environ, 2020, 28, 458–470.

34 M. Tenkanen, T. Hausalo, M. Siika-aho, J. Buchert and L. Viikari, in The 8th International Symposium on Wood and Pulping Chemistry, Gummerus, Finland, 1995, vol. 3, pp. 189–194.

35 J. B. Sumner and C. V. Noback, Journal of Biological Chemistry, 1924, 62, 287–290.

36 M. C. Flickinger, J. L. Schottel, D. R. Bond, A. Aksan and L. E. Scriven, Biotechnology Progress, 2007, 23, 2–17.

37 C. F. Meunier, P. Dandoy and B.-L. Su, Journal of colloid and interface science, 2010, 342, 211–224.

38 X. Pu, Y. Wu, J. Liu and B. Wu, Chem Bio Eng., 2024, 1, 568–592.

39 S. Vajravel, S. Sirin, S. Kosourov and Y. Allahverdiyeva, Green Chemistry, 2020, 22, 6404–6414.

40 C. Nagy, K. Thiel, E. Mulaku, H. Mustila, P. Tamagnini, E.-M. Aro, C. C. Pacheco and P. Kallio, Microb Cell Fact, 2021, 20, 130.

41 Carlos Eduardo De Farias Silva, Barbara Gris, Eleonora Sforza, Nicoletta La Rocca, and Alberto Bertucco, Chemical Engineering Transactions, 2016, 49, 241–246.

42 E. Rueda, A. Álvarez-González, J. Vila, R. Díez-Montero, M. Grifoll and J. García, Science of The Total Environment, 2022, 829, 154691.

43 J. Kurkela and T. Tyystjärvi, Physiologia Plantarum, 2024, 176, e14140.

44 Y. Ohashi, W. Shi, N. Takatani, M. Aichi, S. Maeda, S. Watanabe, H. Yoshikawa and T. Omata, Journal of Experimental Botany, 2011, 62, 1411–1424.

45 T. V. Laurinavichene, S. N. Kosourov, M. L. Ghirardi, M. Seibert and A. A. Tsygankov, Journal of biotechnology, 2008, 134, 275–7.

46 D. J. Dickson and R. L. Ely, Appl Microbiol Biotechnol, 2013, 97, 1809–1819.

47 F. Khosravitabar and C. Spetea, International Journal of Hydrogen Energy, 2024, 67, 925–932.

48 K. R. Sawant, P. Savvashe, D. Pal, A. Sarnaik, A. Lali and R. Pandit, Bioresource Technology, 2021, 341, 125852.

49 I. Homma, T. Isogai, T. Saito and A. Isogai, Cellulose, 2013, 20, 795–805.

50 B. P. Frank, C. Smith, E. R. Caudill, R. S. Lankone, K. Carlin, S. Benware, J. A. Pedersen and D. H. Fairbrother, Environ. Sci. Technol., 2021, 55, 10744–10757.

51 D. Gökçe, M. D. Şeftalicioğlu, B. A. Erden and S. Köytepe, Water Air Soil Pollut, 2022, 233, 434.

52 Y. Ding, Y. Dong, M. Ma, L. Luo, X. Wang, B. Fang, Y. Li, L. Liu and F. Ren, iScience, 2023, 26, 108434.

53 M. Tayyab, M. Dreis, D. Blaudszun, K. Pellumbi, U. Nzotcha, H. Tempel, M. Q. Masood, H. Weinrich, S. Stießel, K. J. Puring, R.-A. Eichel and U.-P. Apfel, Energy Environ. Sci., 2025, 10.1039.D4EE06204C.

54 L. Zeng, Y. Cao, Z. Li, Y. Dai, Y. Wang, B. An, J. Zhang, H. Li, Y. Zhou, W. Lin and C. Wang, ACS Cataysis, 2021, 11, 11696–11705.

55 J. Ding, W. Zhao, L. Zi, X. Xu, Q. Liu, Q. Zhong and Y. Xu, International Journal of Hydrogen Energy, 2020, 45, 15254–15262.

