## Supplemental Figure 1 for "Over four months of ethylene production: Unlocking the potential of solid-state photosynthetic cell factories"

**Suppl. Fig. 1** Validation of *efe* gene integrity during long-term cultivation of ethylene-producing *Synechocystis* cells entrapped within thin nanocellulose films compared to suspension culture.

Representative segment of the alignment:

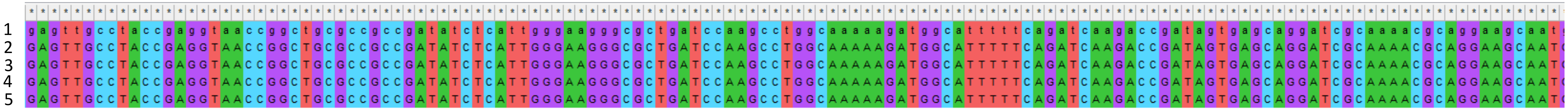


Full sequence alignment:

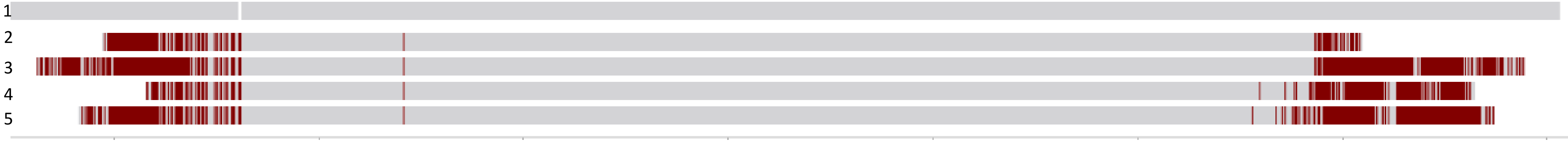


Sequencing analysis of the *efe* gene from *Pseudomonas syringae* confirms the genetic stability of the plasmid in various cultivation conditions and at different time points. The reference sequence (#1) corresponds to the *efe* gene from *P. syringae*. Experimental samples were collected from:

- #2 the film at the beginning of the experiment;
- #3 the film after 142 days of cultivation (see Fig. 1B in the main paper);
- #4 the film after 43 days of cultivation with 120 mM bicarbonate supplementation (see Fig. 2A);
- #5 the PhBFR with suspension culture at the end of the experiment (see Fig. 3).
